# A gene expression atlas of a *bicoid*-depleted *Drosophila* embryo reveals early canalization of cell fate

**DOI:** 10.1101/004788

**Authors:** Max V. Staller, Charless C. Fowlkes, Meghan D.J. Bragdon, Zeba B. Wunderlich, Angela H. DePace

## Abstract

In developing embryos, gene regulatory networks canalize cells towards discrete terminal fates. We studied the behavior of the anterior-posterior segmentation network in *Drosophila melanogaster* embryos depleted of a key maternal input, *bicoid (bcd),* by building a cellular-resolution gene expression atlas containing measurements of 12 core patterning genes over 6 time points in early development. With this atlas, we determine the precise perturbation each cell experiences, relative to wild type, and observe how these cells assume cell fates in the perturbed embryo. The first zygotic layer of the network, consisting of the gap and terminal genes, is highly robust to perturbation: all combinations of transcription factor expression found in *bcd* depleted embryos were also found in wild type embryos, suggesting that no new cell fates were created even at this very early stage. All of the gap gene expression patterns in the trunk expand by different amounts, a feature that we were unable to explain using two simple models of the effect of *bcd* depletion. In the second layer of the network, depletion of *bcd* led to an excess of cells expressing both *even skipped* and *fushi tarazu* early in the blastoderm stage, but by gastrulation this overlap resolved into mutually exclusive stripes. Thus, following depletion of *bcd,* individual cells rapidly canalize towards normal cell fates in both layers of this gene regulatory network. Our gene expression atlas provides a high resolution picture of a classic perturbation and will enable further modeling of canalization in this transcriptional network.

## Introduction

Specialized cell fates underlie the diversity of metazoan form and function. Cell fates are discrete and intermediate fates are confined to pathologies. These observations led Conrad Waddington to propose that development canalizes cell fate (Waddington, 1942). It remains a grand challenge to unravel how developmental gene regulatory networks robustly differentiate multi-potent progenitors into specialized cells in the face of extensive genetic variability and environmental insults. Cell fates are determined by gene expression patterns, so canalization can be studied using gene expression profiles in individual cells.

The *Drosophila melanogaster* blastoderm embryo is a powerful system for studying the properties of developmental gene regulatory networks. Anterior-posterior patterning is controlled by a well-characterized transcriptional network, where all the key genes and connections are known (Lawrence, 1992; St Johnston and Nüsslein-Volhard, 1992; Jaeger et al., 2013). This network can be perturbed using genetic techniques (St Johnston and Nüsslein-Volhard, 1992; Jaeger et al., 2013), and mRNA or protein levels can be quantitatively measured in every cell of the embryo using imaging (Keränen et al., 2006; Luengo Hendriks et al., 2006; Luengo Hendriks et al., 2007; Pisarev et al., 2009). Images of individual embryos can be combined into a gene expression atlas, which captures the average level of expression for an arbitrary number of genes in the same morphological framework (Fowlkes et al., 2008). Gene expression atlases are well suited for studying canalization because they include data for many genes at cellular resolution. Here, we built an atlas of an embryo depleted of a key transcriptional regulator.

We depleted the maternally deposited transcription factor, *bicoid (bcd). bcd* is at the top of the anterior-posterior patterning network. *bcd* specifies head cell fates by activating many genes and represses tail fates by inhibiting translation of maternal *caudal (cad)* (Driever et al., 1990, Lawrence, 1992; St Johnston and Nüsslein-Volhard, 1992; Dubnau and Struhl, 1996). Embryos laid by *bcd* mutant females lack all head structures and instead have a second set of tail structures (Frohnhöfer and Nüsslein-Volhard, 1986); changes in early zygotic expression patterns of the gap and terminal genes prefigure the conversion of the head into a second tail (Nüsslein-Volhard et al., 1987; Tautz, 1988; Hulskamp et al., 1990; Kraut and Levine, 1991b; Pignoni et al., 1992). This dramatic re-patterning provides a strong perturbation for exploring canalization of cell fate: new cell fates might be produced in these embryos, either transiently or permanently.

Cytoplasmic transplantation experiments suggest that cells re-pattern to existing cell fates (Nüsslein-Volhard et al., 1987). However, the *bcd* mutant cuticle is much smaller than a WT cuticle, raising the possibility that misspecified cells may die by apoptosis later in development (Werz et al., 2005). Using out cellular resolution gene expression atlas of a *bcd*-depleted embryo, we asked whether canalization of cell fate occurs during the blastoderm stage and whether it is controlled by the early segmentation network.

The *bcd*-depleted gene expression atlas we report here combines data for twelve genes and six enhancer reporters into a single morphological framework for 6 time points in stage 5 (blastoderm). This is the first cellular resolution data set for a genetic perturbation that includes all the cells in the embryo. To date, there have been four cellular resolution datasets of zygotic mutants that measured fewer genes in part of the embryo (Janssens et al., 2013; Surkova et al., 2013). To circumvent the difficulty collecting mutant embryos, we used shRNA knockdown (Staller et al., 2013).

Analysis of our atlas data revealed multiple properties of the segmentation network under perturbation. First, after *bcd* depletion, the posterior gap gene expression patterns expand asymmetrically: these patterns expand to fill 75% of the *bcd*-depleted embryo compared to 55% of the WT embryo. Notably, each pattern expanded by a different amount. Second, to measure the degree of canalization, we examined gene expression profiles in individual cells and found that all combinations of genes present in the *bcd*-depleted atlas were also present in WT. Thus, no novel cell fates were identified in the *bcd*-depleted embryo. Finally, we directly observed canalization of the parasegment boundaries during the blastoderm stage: extensive early overlap of *even skipped (eve)* and *fushi tarazu (ftz)* mRNA patterns in *bcd*-depleted embryos resolved into mutually exclusive domains. The *bcd* RNAi gene expression atlas thus highlights two ways in which cell fate specification is canalized in early embryos: 1) new combinations of gap and terminal genes do not appear, and 2) after some delay, the network resolves the early overlap of the *eve* and *ftz* mRNA patterns to establish sharp parasegment boundaries.

## Results

### Maternal Gal4 shRNA knockdown of *bcd* phenocopies mutant alleles

To collect the large quantities of embryos necessary to build a gene expression atlas, we used the “maternal Gal4 shRNA” system to deplete *bcd* mRNA in the female germ line (Ni et al., 2011; Staller et al., 2013). We crossed *maternal triple driver Gal4 (MTD*-*Gal4)* females with *UAS-shRNA-bcd* males and collected embryos laid by *MTD-Gal4/UAS-shRNA-bcd* females (Fig. 1A). shRNAs provide an alternative to sickly mutant stocks because flies have normal fertility and all the embryos have the desired genotype. These are significant advantages for atlas building, which requires large numbers of embryos.

**Fig. 1:**
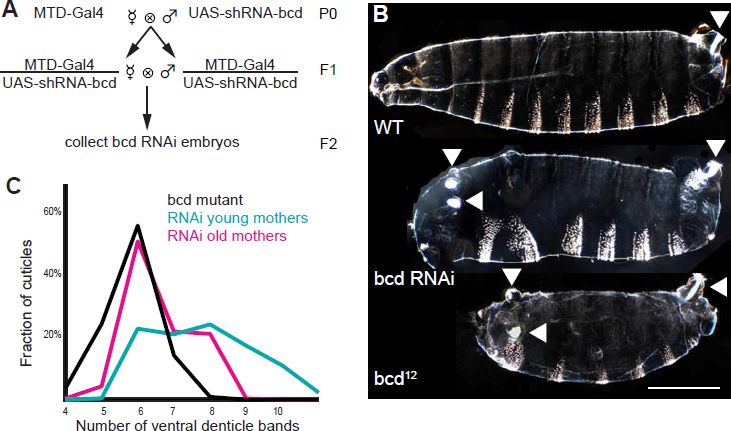
Gal4-driven shRNA against *bcd* in the female germline phenocopies *bcd* mutant alleles A. Crossing scheme for generating *bcd* RNAi embryos (methods). B.Top: dark field image of a WT unhatched larval cuticle with the filzkörper (tail structure) indicated with an arrowhead. The white patches of bristles on each segment are the ventral denticle bands that the larva uses to grip surfaces as it crawls. Middle: *bcd* RNAi cuticle. Bottom: *bcd* mutant cuticle. The *bcd* RNAi embryo has several key features of a classic *bcd* mutant, including the the absence of all head and thoracic structures, and the unextended ectopic filzkörper (arrowheads). All cuticles are oriented with anterior to the left and ventral on the bottom. Scale bar 200 um. C. The strength of knockdown increases and the phenotypic variability decreases as the *MTD-Gal4/UAS-shRNA-bcd* mothers age. Embryos from young and old mothers were collected on day 3 and day 15 respectively. Mutant n = 216, old mothers n = 253, young mothers n = 217. Coefficients of variation (CVs) are: mutant = .1271, old mothers = . 1356, and young mothers = .1800. Representative images of individuals with 4-11 denticle bands are shown in Fig. S1.

Embryos laid by *MTD-Gal4/UAS-shRNA-bcd* females (*bcd* RNAi embryos) have a distribution of weak to strong *bcd* phenotypes that overlaps the distribution of embryos laid by *bcd* mutant females (*bcd* mutant embryos). Larval cuticles with strong *bcd* phenotypes contain only abdominal segments while cuticles with weak *bcd* phenotypes lack head structures and contain a variable number of thoracic segments (Fig. 1B, Fig. S1) (Frohnhöfer and Nüsslein-Volhard, 1986). To compare phenotypic variability in *bcd* RNAi and *bcd* mutant embryos, we counted the number of ventral denticle bands visible on each cuticle (Fig. 1C). To permit alignment and averaging of embryos for atlas building, we sought to reduce the variability of the *bcd* RNAi embryo population as much as possible. The primary determinant of phenotypic strength and variability in *bcd* RNAi embryos was the age of the mothers. Young mothers laid embryos that resemble both weak and strong *bcd* alleles, while older mothers laid embryos with stronger and less variable phenotypes (Fig. 1C). This improvement may stem from a slowing of oogenesis in older females, permitting the shRNAs more time to deplete targets (Ni et al., 2011). The distribution of segment number shifts slowly as the mothers age, as shown by a time course of cuticles collections (Fig. S2A). By day 8, as measured by qPCR, the levels of *bcd* transcripts in 0-2 hr old embryos were <10% of those in embryos laid by *MTD-Gal4/UAS-shRNA-bcd* females (Fig. S2B). To balance the need to reduce variability against declining fecundity, we elected to collect embryos after aging the flies for 11-15 days, at which point >90% of embryos passed our threshold for a strong *bcd* phenotype: having 8 or fewer denticle bands (all of which had abdominal character) and ectopic tail structures.

We investigated several additional potential contributors to variability, including temperature, shRNA sequence, maternal driver and the number of *UAS-shRNA-bcd* transgenes in each embryo, but none contributed strongly (Fig. S2D). The male genotype also had no effect: crossing *MTD-GAL4/UAS-shRNA-bcd* females to various male genotypes had no measurable effect on the cuticle phenotype, confirming that *bcd* RNAi depletion, like the *bcd* mutant, has a purely maternal effect (Fig. S2C) (Frohnhöfer and Nüsslein-Volhard, 1986). The absence of any paternal or zygotic effects enabled us to introduce enhancer *lacZ* reporters into the perturbed embryo by crossing *MTD-Gal4/UAS-shRNA-bcd* females with males homozygous for the reporter (methods; Table S4). We integrated these enhancer reporter expression patterns into the gene expression atlas by co-staining with a fiduciary marker (methods).

### Building a gene expression atlas of a *bicoid* depleted embryo

To build a gene expression atlas, we segment image stacks of individual embryos and use a fiduciary marker gene to align embryos together; this process creates an average picture. Building the *bcd* RNAi gene expression atlas required a morphological template for the position of every cell and a registration template for fine-scale alignment (Fowlkes et al., 2008). To accommodate the small increase in average cell number and changes in cell density patterns in *bcd* RNAi embryos (Fig. S3A,B), we built a new morphological template (methods). In WT, our registration template uses the expression pattern of a pair-rule gene, *eve* or *ftz,* to finely align embryos (Fowlkes et al., 2008). In *bcd* RNAi embryos, both *eve* and *ftz* patterns are severely disrupted, precluding the use of the WT registration template. To create a *bcd* RNAi registration template, we imaged a set of 249 embryos laid by older mothers stained for *ftz* mRNA (Fowlkes et al., 2008; Fowlkes et al., 2011). At late time points, some embryos expressed an extra *ftz* or *eve* stripe (discussed below); we manually excluded these individuals from the atlas (methods). With the new morphological template and registration template of the perturbed *ftz* pattern, we built the *bcd* RNAi gene expression atlas (methods).

The final atlas includes 1567 embryos with mRNA stains for all of the following genes: *caudal (cad), Kruppel (Kr), knirps (kni), giant (gt), hunchback (hb), fork head (fkh), huckebein (hkb), tailless (tll), Dichaete (D), runt (run), even-skipped (eve), fushi-tarazu (ftz)*, distributed over 6 time points in the blastoderm stage (Fig. 2; embryos per gene in Table S1). In addition, we measured 6 lacZ reporters containing the following enhancers: *hb posterior, gt posterior, eve stripe3+7, eve stripe5*, and two *eve stripe4+6* enhancers (Table S4). We collected embryos with the eve stripe1, *eve stripe2*, and *eve late seven stripe* enhancers, but these sequences drove very little expression (discussed below). Finally, we collected protein data for Hb, for which there is a large difference in the mRNA and protein patterns in both WT and *bcd* mutants (Fig. 2) (Tautz, 1988; Struhl et al, 1989; Fowlkes et al., 2008). We divide the data into six temporal cohorts of ∼10 minutes spanning stage 5 (methods).

**Fig. 2:**
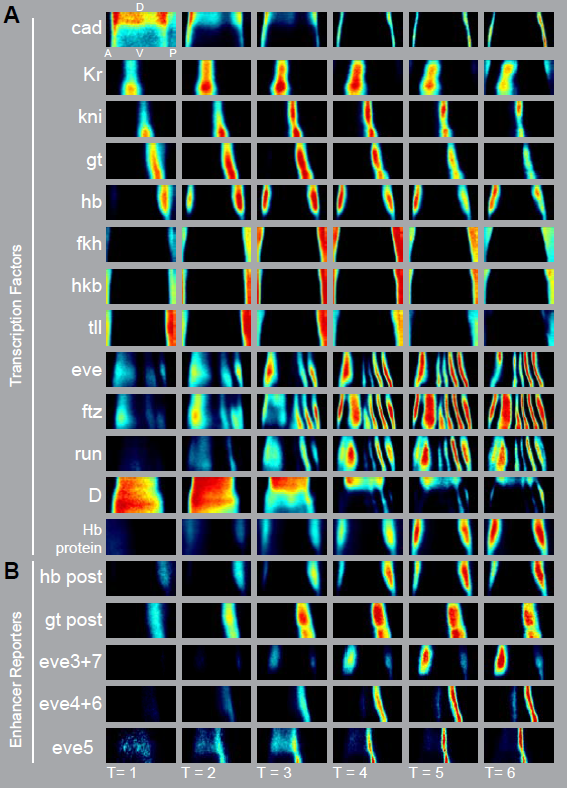
The gene expression atlas of a *bcd* depleted embryo highlights the expansion of trunk patterns, duplication of posterior patterns, and loss of anterior patterns A. Heat maps for mRNA expression patterns of the twelve core members of the segmentation network included in the *bcd* RNAi atlas. Relative mRNA levels scale from no expression (black) to peak expression (red). We also collected protein data for Hb. We partition the data into 6 ∼10 min cohorts that span all of stage 5 using a morphological marker (methods). B. Heat maps of five enhancer lacZ reporter constructs included in the atlas. Anterior is left, dorsal is top.

The *bcd* RNAi gene expression atlas is of similar or higher quality to than the original WT atlas based on two metrics. First, the standard deviation of each gene averaged over all cells and all time points, was smaller in the *bcd* RNAi atlas than in the WT atlas for 10 of the 12 genes (Table S2). Second, for all but a few genes, background expression levels in OFF cells were lower in *bcd* RNAi, as shown in the histogram of expression levels (Fig. 5A).

### Characteristics of the *bcd* RNAi gene expression atlas

The final *bcd* RNAi gene expression atlas combines data for twelve genes and six reporters into a single morphological framework for the entire embryo at cellular resolution. These patterns qualitatively agree with published images (Frasch and Levine, 1987; Nüsslein-Volhard et al., 1987; Tautz, 1988; Struhl et al., 1989; Hulskamp et al., 1990; Kraut and Levine, 1991b; Rivera-Pomar et al., 1995). Key patterning genes in the trunk are still expressed, but positions and levels change. Consistent with the *bcd* mutant literature, all gap gene expression domains in the anterior 45% of WT were lost and replaced with a second set of posterior expression patterns for *tll, hkb, fkh* and *hb* (Tautz, 1988; Pignoni et al., 1992). The improved temporal resolution of our data revealed that the ectopic *hb* and *fkh* patterns are delayed compared to the corresponding patterns in the tail (Fig. 2). The posterior gap patterns moved anteriorly over time as in WT (Jaeger et al., 2004b); in a mirror image fashion, the ectopic *hb* pattern moved away from the anterior pole (Figs 2, 4A). All measured pair-rule patterns were perturbed: at the end of the cellular blastoderm stage, *bcd* RNAi embryos have 5 *eve* stripes, 6 *ftz* stripes, and 6 *runt* stripes instead of the normal 7 stripes.

There is more variability in the pair-rule gene expression patterns than in the gap gene expression patterns. In 22% (22/98) of *bcd* RNAi embryos the anterior *eve* stripe split in two at T=5 and T=6. These embryos were excluded from the atlas, as described above. In embryos with a single anterior stripe, the position and width of this stripe varied more than the other stripes (Fig. S4). The boundaries of both the reporters and endogenous *eve* stripes refine later in *bcd* RNAi than in WT (Fig. 2; Fig. 3). Stripes 6 and 7 start out wide, but by gastrulation narrow to be only slightly wider than WT (Fig. S4). Aside from the anterior stipe, eve stripe width coefficients of variation (CVs) were comparable to gap gene width CVs, consistent with each layer of the network having similar amounts of variability (Fig. S5).

**Fig. 3:**
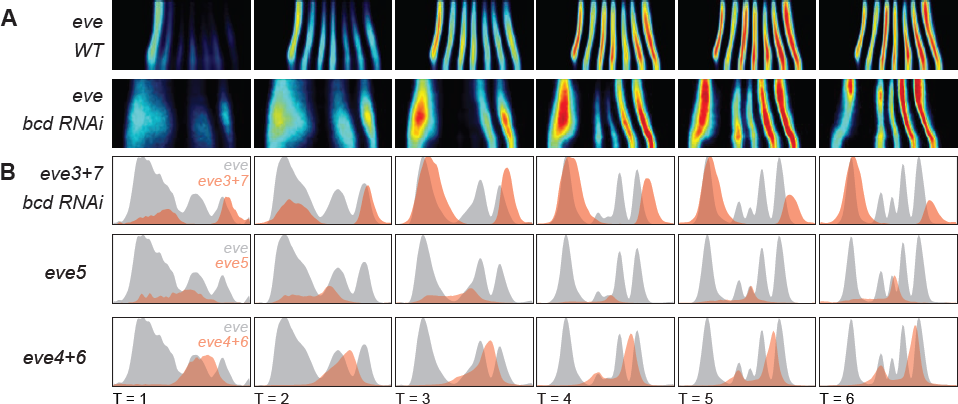
Enhancer reporter constructs identify all of the *eve* stripes in *bcd* RNAi embryos A. Heat maps for *eve* mRNA patterns in the WT and *bcd* RNAi gene expression atlases. B. Line traces of the endogenous *eve* pattern (gray) and the enhancer reporter pattern (orange) plot anterior-posterior position on the x-axis and expression level on the y-axis for a single strip along the side of the embryo. The levels of the enhancer reporter line traces have been manually scaled to match the corresponding endogenous stripe.

### Identifying the perturbed *eve* stripes in *bcd* RNAi embryos

To correspond the five *eve* stripes in *bcd* RNAi embryos with their WT counterparts, we introduced *eve* enhancer reporter constructs into the *bcd* RNAi background (Fig. 3). The *eve* locus contains five enhancers that together drive seven stripes: stripes 1, 2 and 5 each have a dedicated enhancer, while stripes 4+6 and stripes 3+7 are produced as pairs from two different enhancers (Goto et al., 1989; Small et al., 1991; Small et al., 1996; Fujioka et al., 1999). In *bcd* mutants, reporter constructs carrying the *eve* stripe 2 (*eve2),* and *eve* stripe 1 (*eve1)* enhancers are not expressed, while the *eve* stripe 3+7 enhancer (*eve3+7*) drives an anteriorly shifted and expanded stripe 3 and a reduced stripe 7 (Small et al., 1991; Small et al., 1996; Ochoa-Espinosa et al., 2005). To our knowledge, the stripe 4+6 and stripe 5 enhancer reporter constructs have not previously been examined in *bcd* mutant embryos.

We found that the five *eve* stripes in *bcd* RNAi embryos correspond to *eve* stripes 3+7, *eve* stripes 4+6 and *eve* stripe 5 (Fig. 3). The *eve2* reporter showed some activity in a handful of *bcd* RNAi embryos, but since our version of this reporter drives some expression of stripe 7 in WT (Goto et al., 1989; Small et al., 1991), it is likely that this residual expression is a duplication of the stripe 7 activity in the anterior (Fig. S9). Together, the reporters qualitatively account for all of the *eve* stripes in *bcd* RNAi embryos, although the agreement for stripes 6 and 7 is notably worse. We are quantifying and investigating these discrepancies in ongoing work.

### Two simple models fail to predict the asymmetric expansion of gap gene expression patterns

The most prominent feature of the *bcd* RNAi embryo is the asymmetric expansion of the posterior gap gene expression patterns. In WT, the anterior boundary of Kr begins at 44% egg length (from the anterior) and the *Kr, kni, gt, hb* and *tll* patterns fill the remaining 55% of the embryo. In *bcd* RNAi, the anterior boundary of *Kr* shifts to begin at 27% egg length (from the anterior), the four gap gene domains expand to fill 73% of the embryo (Fig. 4). While individual pattern shifts have been noted in the past (Struhl et al., 1989; Hulskamp et al., 1990; Kraut and Levine, 1991b; Rivera-Pomar et al., 1995), our measurements revealed that at early time points each pattern expanded by a different amount (Fig. 4B). Each domain contracted or expanded with unique dynamics, and at the last time point *kni* was narrower than in WT (Fig. 4B).

**Fig. 4:**
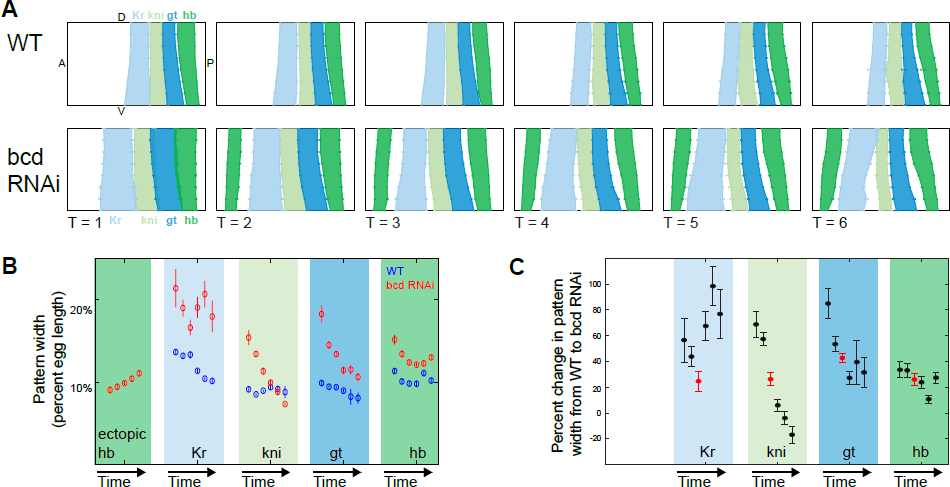
The gap gene expression patterns in the trunk expand by different amounts in *bcd* RNAi embryos A. The gap gene expression patterns in the trunk in WT and *bcd* RNAi gene expression atlases plotted as unrolled embryos. The pattern boundaries were calculated by finding the inflection point of the pattern in individual embryos. Error bars are standard error of the mean. B. The widths of each gap gene expression domain change over time in WT (blue) and *bcd* RNAi (red). For each gene, the width of the pattern, calculated from a lateral strip, is plotted over 6 time points. The patterns narrow over time in both genotypes, but more quickly in *bcd* RNAi. Pattern widths are shown in percent of egg length. For reference, a nuclear diameter is about 1% of egg length. C. The percent change in gap gene expression domain widths between WT and *bcd* RNAi, calculated for each time point from C. Time point 3 is indicated in red.

**Fig. 5:**
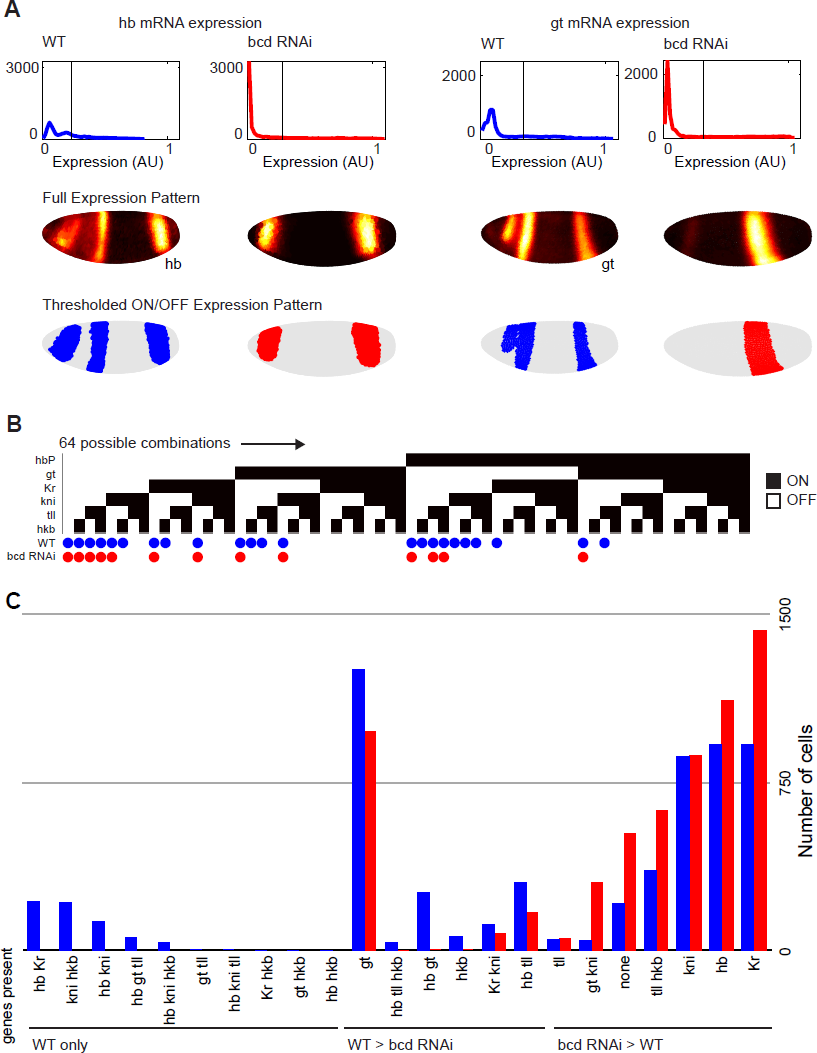
There are no new combinations of gap and terminal gene expression patterns in *bcd* RNAi embryos. A. For each gene, we threshold the expression pattern to find ON cells. Histograms of expression levels (top), heat maps of continuous expression patterns (middle), and the thresholded pattern (bottom) are shown for two genes in WT and *bcd* RNAi. B. Each column represents one of the 64 possible ON/OFF combinations of 6 genes. Black boxes indicate presence of a transcription factor in a combination. There are 23 combinations present in WT (indicated by blue dots) and 13 combinations present in *bcd* RNAi (red dots). No combinations are present only in *bcd* RNAi. C. The number of cells with each combination in each genotype. Virtually all combinations change in abundance. In WT, there are two regions of cells expressing only *gt* and in *bcd* RNAi there is only the expanded posterior region. Similarly, for *kni* in WT there are two regions expression only *kni* and one expanded posterior region in *bcd* RNAi.

We tested two phenomenological models for how the gap patterns expand in the *bcd* RNAi embryo: linear scaling of the whole pattern and boundary positioning by an exponentially distributed morphogen gradient. First, we tested a model in which the entire pattern stretches to fill the space and each individual pattern expands by the same percentage. Recalculating the expansions as percentages shows that this model almost matches the data at T=3, but diverges from the data at other times (Fig. 4C).

Second, we considered a model where an exponential gradient of Bcd protein sets boundaries in a concentration-dependent manner (Alon, 2007; Barkai and Shilo, 2009) (reproduced in Supplemental Note 1). There is evidence in the literature that Bcd protein levels directly set the boundaries of *Kr* and *kni* (Jacob et al., 1991; Rivera-Pomar et al., 1995). This model predicts that each boundary should shift by the same amount (Fig. S9). However, this model failed to match the data at any time point, consistent with evidence that these patterns are also regulated by maternally deposited *hb* and *cad* (Hulskamp et al., 1990; Rivera-Pomar et al., 1995). The asymmetric expansion of the gap genes is an important feature of our dataset that can be used to challenge other models of gap gene pattern formation and refinement (Jaeger et al., 2004a; Jaeger et al., 2004b; Bieler et al., 2011; Hengenius et al., 2011; Papatsenko and Levine, 2011).

### Gap gene expression patterns are canalized in *bcd* RNAi embryos

Depletion of *bcd* causes a dramatic perturbation of the organismal body plan, leading to complete replacement of the head segments with a second set of tail structures. The fact that the head region largely phenocopies the tail region suggests that there is a strong canalization of cell fate from anterior to posterior (Nüsslein-Volhard et al., 1987). Since our gene expression atlas measures the activity of the core patterning network in all cells of the embryo, we can assess the degree of this canalization at the molecular and cellular level. For example, do any cells in the *bcd* RNAi embryo fail to reach a cell state that is recognizable in the WT embryo? That is, are any new cell fates created?

We use a simplified definition of a cell fate: a cell fate corresponds to a binary gene expression profile, where each gene is either ON or OFF. The emergence of a new combination of genes in *bcd* RNAi embryos would indicate a qualitatively new cell fate, although the absence of new combinations by this definition does not rule out more subtle changes in cell identity. We analyzed combinations of genes in the first zygotic layer of the network: the gap and terminal genes *Kr, hb, gt, kni, tll* and *hkb*. We used mRNA patterns in our third time point as a snapshot of cell fates. Blastoderm cell fates are transient, not terminal fates, but using gene expression allows us to study how they are specified. For each of the 6 regulators, we thresholded expression to classify cells as ON or OFF (Fig. 5A; methods; Table S3), giving 2^6^ (64) possible ON/OFF combinations. There are three possible outcomes: a cell fate can be present only in WT, present only in *bcd* RNAi, or present in both. Our analysis implicitly assumes that all six genes contribute to cell fate, a reasonable assumption for anterior-posterior patterning.

Of the possible 64 combinations, 23 cell fates were present in WT embryos (Fig. 5B). In WT, there were no combinations with 4 or more genes and only 4/20 possible combinations of 3 genes, consistent with the strong mutual repression between some of the gap genes (Jaeger, 2011). All cells in *bcd* RNAi embryos fell into 13 cell fates, all of which were present in WT; the 10 WT cell fates that were lost in *bcd* RNAi had most cells located in the head (Fig. S6). Virtually all the shared cell fates changed in abundance between genotypes, with 6 more abundant in WT, and 7 more abundant in *bcd* RNAi (Fig. 5C). By this simplified definition, no new cell fates were created following *bcd* depletion, but existing cell types change in proportion. This result suggests that the first zygotic layer of the gene regulatory network canalizes cells towards normal fates.

Our conclusions are supported even when we modulated the ON/OFF threshold for gene expression.. For T=3, we found that all combinations of genes in *bcd* RNAi were also present in WT even after modulating the threshold (Fig. S7). At other times and thresholds, we sometimes found a handful of cells with a combination unique to *bcd* RNAi. In virtually all cases, however, we also found the combination in WT at other thresholds or adjacent time points. This finding suggested that these cases are not true new cell fates, but instead stem from the fact that each gene is normalized separately in each genotype. One advantage of thresholding gene expression patterns in the combination analysis is that it mitigates the effect of the differences in distribution of OFF cell expression levels between the atlases. This difference likely comes from improvements in the staining protocol. Each gene is normalized separately in each atlas and the WT data tend to have higher background levels (Fig. 5A). These differences confound many analyses, including clustering and the expression distance metric previously used to compare species (Fowlkes et al., 2011). There are a few additional cases that may be due to changes in expression dynamics or differences in data quality. At T=6 there were a handful of cells with new combinations of the terminal patterns of *tll*, *hkb*, and *Kr* in *bcd* RNAi (Fig. S7). These new combinations may reflect changes in the dynamics of *tll* expression or the low quality of the WT T=6 *hkb* data (Fig. S7C). When we substituted Hb protein data for *hb* mRNA data, we again found that no new combinations arose in *bcd* RNAi, but for a more limited range of thresholds, consistent with our empirical finding that the Hb protein data is harder to faithfully partition into ON and OFF cells.

### The pair-rule gene expression patterns of *eve* and *ftz* are dynamically canalized in *bcd* RNAi embryos

The primary pair-rule genes *eve* and *ftz* define the parasegment boundaries that later establish the compartment boundaries (Martinez-Arias and Lawrence, 1985; Lawrence, 1992). We chose to examine this layer of the network separately from the gap and terminal genes for three reasons: 1) in addition to being regulated by the gap genes, *eve* and *ftz* take direct input from the maternal genes; 2) these genes may be sensitive to quantitative changes in relative levels of the gap genes that are not revealed by our binary combination analysis; and 3) while the initial gap gene patterns appear in stage 4, before we started collecting data, our stage 5 data capture the emergence, establishment, and refinement of *eve* and *ftz* expression. In WT, these two gene expression patterns are mutually exclusive for virtually the entire blastoderm stage (Fig. 6). In *bcd* RNAi, some individual embryos had extensive overlap of these two patterns. To quantify this difference, we examined individual embryos stained for *eve* and *ftz*, thresholded expression to assign ON/OFF states for each gene, and counted the fraction of cells expressing both. In WT <10% of cells express both genes in the first time point and this fraction quickly drops. In contrast, in *bcd* RNAi embryos at T=1 and T=2, ∼20% of cells express both *eve* and *ftz.* Beginning at time point 3, the fraction of cells in *bcd* RNAi embryos expressing both genes drops sharply (Fig 6B). The shape of the trend does not depend on the threshold used to assign cells as ON and OFF (Fig. S8). In our dataset, early *eve/ftz* overlap resolves into mutually exclusive stripes, another manifestation of canalization in the segmentation network.

**Fig. 6:**
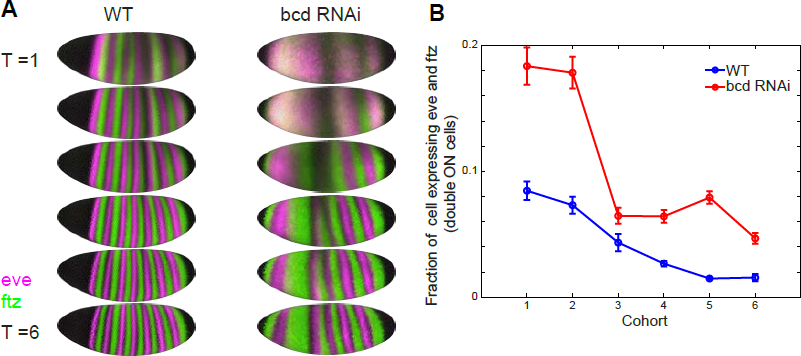
The mRNA expression patterns of *eve* and *ftz* canalize over the blastoderm stage A. *eve* (pink) and *ftz* (green) mRNA patterns in the gene expression atlas for each cohort in WT and *bcd* RNAi. Cells without any expression appear black and cells expressing both *eve* and ftz appear white. B. We quantified the fraction of cells that express both *eve* and *ftz* in individual embryos. For each embryo, we thresholded each expression pattern to be ON or OFF, and counted the fraction of cells where both genes were ON. WT N = 113, *bcd* RNAi N = 287.

## Discussion

We used the maternal Gal4 shRNA system (Staller et al., 2013) to build a gene expression atlas of an embryo depleted of *bcd*, a maternally deposited transcription factor critical for anterior-posterior patterning of the *Drosophila melanogaster* embryo. This atlas captures multiple gene expression patterns after a genetic perturbation, in 3D, at cellular resolution, for the whole embryo, for the first time. In the absence of *bcd,* the segmentation network canalizes cell fates: new combinations of gap and terminal genes are not expressed after *bcd* depletion. The widths of the gap gene expression patterns expanded asymmetrically in a way that cannot be explained by simple re-scaling, or by assuming that Bcd protein levels directly set boundary positions. Downstream of the gap and terminal system, the expression patterns of pair-rule genes *eve* and *ftz* overlap in more cells and for more time in *bcd* RNAi embryos compared to WT embryos, but eventually establish sharp parasegment boundaries. We conclude that the anterior-posterior patterning network robustly specifies cell fates following the loss of a key maternal input.

### Phenotypic variability in *bcd* RNAi embryos can be controlled, and may be useful in the future

To build the gene expression atlas, we suppressed the variability in the distribution of phenotypes in *bcd* RNAi embryos by using specific collection conditions and manually curating the dataset. In the future, such variability may be useful for studying other emergent properties of the network. For example, partially penetrant mutants helped constrain mathematical models of signal integration (Corson and Siggia, 2012). Some of our phenotypic variability likely stems from inconsistent shRNA knockdown (Mohr and Perrimon, 2012), but some undoubtedly results from the response of the network to the loss of *bcd* (Fig. 1C). Increased variability in mutant phenotypes is common (Waddington, 1942; Wieschaus et al., 1984), and recent examination of gene expression patterns in *tll, Kr, kni,* and *Kr/kni* mutants concluded that there was more molecular variability in mutant embryos than in WT embryos (Janssens et al., 2013; Surkova et al., 2013).

One especially dramatic example of variability in *bcd* RNAi embryos was the splitting of the anterior *eve* stripe in 22% of embryos at T=5 and T=6. Since *eve* and *ftz* expression domains define the parasegment boundaries (Martinez-Arias and Lawrence, 1985), this molecular variability in the blastoderm may explain the variable number of denticle bands we see in *bcd* RNAi cuticles. To enable study of this variability, we will make the data from embryos with split anterior *eve* stripes publicly available (depace.med.harvard.edu).

### The segmentation network fills space through asymmetric expansions

The gap gene expression patterns all expand by different amounts in *bcd* RNAi embryos, consistent with the self-organizing behavior of this network (Nüsslein-Volhard et al., 1987; Driever and Nüsslein-Volhard, 1989; Liu et al., 2013). The absence of uniform shifts in boundary positions rules out models where an exponential gradient of *bcd* is primarily responsible for patterning borders (Alon, 2007; Barkai and Shilo, 2009; Ben-Zvi et al., 2011; Tamari and Barkai, 2012). Asymmetric expansions similarly rule out linear scaling models and indicate that a nonlinear mechanism is operating. The gap genes exhibit extensive asymmetric cross-repression (Jäckle et al., 1986; Kraut and Levine, 1991a; Jaeger et al., 2004a; Papatsenko and Levine, 2011), and this cross-repression is stronger than Bcd's instructive role in setting boundaries (Ochoa-Espinosa et al., 2005; Ochoa-Espinosa et al., 2005; Chen et al., 2012). Multiple dynamic models if the gap-gene network have been proposed (Jaeger et al., 2004a; Jaeger et al., 2004b; Bieler et al., 2011; Hengenius et al., 2011; Papatsenko and Levine, 2011), and quantitative data for *Kr* mutant embryos led to refinements in one well-established model of cross-repression (Kozlov et al., 2012). We anticipate that the asymmetric expansion of gap patterns in the *bcd* RNAi embryo will provide a useful challenge for dynamic models investigating this network-level behavior.

### The segmentation network canalizes cell fate in *bcd* RNAi embryos

We found molecular evidence for canalization of cell fate both in the gap gene and pair-rule gene expression patterns. In response to *bcd* depletion, the transcriptional regulatory network does not generate new combinations of gap gene expression patterns—all cells assume fates that already exist in WT embryos. Early overlap of *eve* and *ftz* expression patterns resolves into mutually exclusive stripes. Our analysis is the first direct evidence indicating that the anterior-posterior patterning network prevents the creation of new cell fates in the absence of a maternal input. Several lines of evidence predicted that mutant cell types would eventually adopt wild-type fates, including mutant cuticle phenotypes and cytoplasmic transplantation experiments (Nüsslein-Volhard et al., 1987), the coordinated shifts in blastoderm expression patterns following *bcd* dosage changes (Driever and Nüsslein-Volhard, 1989; Struhl et al., 1989; Liu et al., 2013), the abundant cross repression in the gap gene network (Jäckle et al., 1986; Jaeger et al., 2004b; Papatsenko and Levine, 2011), and the molecular canalization of gene expression patterns in WT (Manu et al., 2009a; Manu et al., 2009b; Gursky et al., 2011). However, it has not been clear when this canalization occurs: immediately, due to the transcriptional network, or later, due to downstream compensatory processes? There is extensive apoptosis in *bcd* mutant embryos later in development, which was proposed to be removal of misspecified cells (Werz et al., 2005). Misspecification can either imply the presence of too many cells of a given type, or the emergence of new types. We have shown that canalization occurs early and strongly, resulting in changes in the abundance of most cell fates, but no cells with new fates.

We propose that the increased apoptosis observed in *bcd* mutants is not pruning cells with new fates, but instead compensating for enlarged compartments. The *eve* and *ftz* stripes set compartment size, and large compartments experience increased cell death while small compartments experience reduced apoptosis (Namba et al., 1997; Hughes and Krause, 2001). The wide second *ftz* stripe (Fig. 2) is approximately where the most apoptosis is observed in *bcd* mutant embryos (Werz et al., 2005). According to our analysis, cells undergoing apoptosis do not have new fates at the blastoderm stage. Rather, they reside in a compartment that is too large, and this increased compartment size may trigger cell death.

### Dynamic canalization establishes sharp *eve* and *ftz* parasegment boundaries

We observed the dynamics of the canalization of parasegment boundaries by examining the expression patterns of *eve* and *ftz* in individual *bcd* RNAi embryos. At T=1 and T=2 ∼20% of cells in *bcd* RNAi express both *eve* and *ftz*, more than double the fraction in WT. By T=3, however, the fraction of cells expressing both genes drops sharply as the patterns resolve into mutually exclusive stripes. Similar early overlaps of *eve* and *ftz* that resolve to mutually exclusive stripes have recently been reported in *Kr* mutant embryos (Surkova et al., 2013). This resolution of *eve* and *ftz* boundaries is likely mediated by direct repression of *ftz* by *eve* and indirect repression of *eve* by *ftz* (Manoukian and Krause, 1992; Saulier-Le Drean et al., 1998; Nasiadka and Krause, 1999; Schroeder et al., 2011). Resolution of compartment boundaries may be a general feature of compensation in mutants, and we hope to uncover the mechanisms and regulatory DNA responsible.

### Conclusion

Reexamining a classic perturbation at cellular resolution provided direct evidence that cell fates are canalized early and strongly by the anterior-posterior network. Our increased resolution also revealed new features of the network, including the asymmetric expansion of the gap genes and the dynamic canalization of the parasegment boundaries. We anticipate that the *bcd* RNAi gene expression atlas will be useful to the modeling community by providing a cellular resolution dataset for testing models of how individual regulatory circuits position expression domains. These studies also lay important groundwork for our long-term goal of identifying the features of the network architecture that contribute to canalization of cell fate. This network has mutual repression in every layer: among the maternal genes, the gap genes, and the pair-rule genes. We speculate that these multiple layers of mutual repression underlie the early and strong canalization of cell fate we observe.

## Materials and Methods

### Fly Work

We depleted *bcd* with *UAS-shRNA-bcd* (TRiP GL00407) and the *maternal triple driver Gal4 (MTD-Gal4)* (Fig. 1A) (Staller et al., 2013). For reference, we used the *bcd^12^* loss of function allele (Bloomington 1755) (Frohnhöfer and Nüsslein-Volhard, 1986). For validation controls we also used the *maternal-tubulin-Gal4 (mat-tub-Gal4)* drivers, GL01320 *UAS-shRNA-bcd*, and TB184 *UAS-shRNA-GFP* (Fig. S2).

To test reporter constructs, we crossed virgin *MTD-Gal4/UAS-shRNA-bcd* females to males homozygous for the reporter gene. Each enhancer was cloned into the *NotI* and *BglII* sites of the *pBOY-lacZ* reporter construct (Hare et al., 2008) and integrated at the attP2 landing pad (Groth et al., 2004). The full list of reporter constructs, sequences, original references and cloning primers are listed in Table S4.

### Preparation of unhatched larval cuticles

Larval cuticles were prepared and mounted in lactic acid (Stern, 2000). We manually counted the number of denticle bands on each cuticle under dark field illumination, rounding up partial segments. For the majority of cuticles shown, a Z-stack of 2-4 images was computationally flattened with Helicon Focus (Helicon soft).

### quantitative RT PCR

Embryos were collected for 2 hours and snap frozen in liquid nitrogen. We extracted RNA with Trizol and synthesized cDNA with superscript reverse transcriptase (Life). We used TaqMan probes (Life) with *actin* as a reference.

### *in situ* hybridization

All RNA stains were performed as in (Fowlkes et al., 2011; Wunderlich et al., 2012). Briefly, embryos were collected over 4 hours at 25°C, dechorionated in bleach, fixed in formaldehyde/heptane for 25 minutes, dehydrated with methanol and stored in ethanol at −20°C. We used a digitonin (DIG) *ftz* probe, a dinitrophenol (DNP) probe against the gene of interest, and developed them sequentially with a tyramide amplification reaction (Perkin Elmer), with DIG in the coumarin channel and DNP in the Cy3 channel. After RNase treatment overnight at 37°C, DNA was stained with Sytox green (Invitrogen). Embryos were dehydrated with ethanol, cleared with xylenes and mounted in DePeX (Electron Microscopy Sciences). To acquire Hb protein data, we stained embryos first with *ftz* DNP in the coumerin channel, and stained with guinea pig anti-Hb (a generous gift from John Reinitz (Chicago, Illinois, USA)) and goat anti-guinea pig AlexaFluor 555 (Life).

### Image acquisition

We acquired Z-stacks with 2-photon excitation at 750 nm, with 1 um increments, and simultaneously collected the 3 fluorescent channels. Protein stains were imaged in the same way. We use automated image processing to segment the nuclei and extract expression of the two genes in every cell, creating a pointcloud file for each embryo (Luengo Hendriks et al., 2006). We manually classified embryos into 6 cohorts: 0-3%, 4-8%, 9-25%, 26-50%, 51-75%, and 76-100% membrane invagination, which evenly divide the ∼60 min blastoderm stage (Keränen et al., 2006).

### Manual data curation

To remove individual embryos with low knockdown from the set of embryos laid by old mothers, we manually inspected the *ftz* pattern. For time points 4-6, we removed embryos with a narrow second *ftz* stripe or an extra *ftz* stripe. For *eve* stains, we removed any embryos with a split anterior stripe.

### Finding expression pattern boundaries

Pointcloud files were manipulated in MatLab (Mathworks) using the pointcloud toolbox (bdtnp.lbl.gov). For each embryo, we created line traces for 16 stripes around the dorsal ventral axis, and found the inflection point in each trace. Similar results were obtained when used the half maximum of each line trace.

### Building the *bcd* RNAi gene expression atlas

To account for a small increase in cell number and changes in cell density, we built a new morphological template for the *bcd* RNAi atlas using 1567 embryos (Fowlkes et al., 2008; Fowlkes et al., 2011). To build a new gene expression registration template we used 249 embryos stained only with DNP *ftz* probes (Fowlkes et al., 2008; Fowlkes et al., 2011). For all other genes, the DIG *ftz* stain in each embryo was used to finely align individual pointcloud files to this registration template. Each gene was normalized separately so that relative levels between time points were preserved, but the absolute levels between atlases are likely different. Cell density maps shown in Fig. S3 were generated using the demo_densities function in the pointcloud toolbox.

Accompanying this paper, we have provided the *bcd* RNAi gene expression atlas and a bundled file containing all the individual embryos stained for *eve* and *ftz*, including those that were excluded from the atlas (depace.med.harvard.edu).

### Identifying combinations of ON and OFF cells

To create the binary gene expression profile of each cell, we thresholded *Kr, hb, kni, gt, tll,* and *hkb* mRNA at T=3. The ON/OFF threshold was calculated for each gene in each atlas, by identifying the mode of the distribution of expression values (the peak of the OFF cell population) and adding one standard deviation. For *eve* and *ftz*, we determined thresholds for each gene in each embryo and recorded the fraction of cells expressing both. Using the published stains of WT embryos, we found that swapping the haptens (DNP and DIG) did not change the fraction of double ON cells (Fig. S8A) (Fowlkes et al., 2008).

## Acknowledgements

We thank John Reinitz for the spectacular Hb antibody, Norbert Perrimon for discussions of phenotypic variability, TRiP for the maternal drivers and initial *UAS-shRNA-bcd* line, Miki Fujioka for providing the *eve late seven stripe* enhancer reporter line, and Tara Lydiard-Martin for making the other *eve* enhancer reporter lines. We thank Ashley Wolf, Becky Ward and members of the DePace Lab for feedback on the manuscript andBen Vincent for extensive comments on the manuscript. This work was supported in part by NSF DBI-1053036 (CCF), the Harvard Herchel Smith Fellowship (MVS) and NIH U01 GM103804-01A1 (AHD).

## Author Contributions

MVS and AHD designed the study. MVS and MDJB performed the experiments. CCF built the templates for the gene expression atlas. ZBW processed raw image stacks into pointcloud files. MVS analyzed the data with input from AHD and ZBW. MVS and AHD wrote the paper.

## References

Alon, U. (2007). Introduction to Systems Biology: And the Design Principles of Biological Networks.

Barkai, N. and Shilo, B. Z. (2009). Robust generation and decoding of morphogen gradients. Cold Spring Harb Perspect Biol 1, a001990.

Ben-Zvi, D., Shilo, B. Z. and Barkai, N. (2011). Scaling of morphogen gradients. Curr Opin Genet Dev 21, 704–710.

Bieler, J., Pozzorini, C. and Naef, F. (2011). Whole-Embryo Modeling of Early Segmentation in Drosophila Identifies Robust and Fragile Expression Domains. Biophysj 101, 287–296.

Chen, H., Xu, Z., Mei, C., Yu, D. and Small, S. (2012). A system of repressor gradients spatially organizes the boundaries of Bicoid-dependent target genes. Cell 149, 618–629.

Corson, F. and Siggia, E. D. (2012). Geometry, epistasis, and developmental patterning. Proc Natl Acad Sci U S A 109, 5568–5575.

Driever, W. and Nüsslein-Volhard, C. (1989). The bicoid protein is a positive regulator of hunchback transcription in the early Drosophila embryo. Nature 337, 138–143.

Driever, W., Siegel, V. and Nusslein-Volhard, C. (1990). Autonomous determination of anterior structures in the early Drosophila embryo by the bicoid morphogen. Development 109, 811–820.

Dubnau, J. and Struhl, G. (1996). RNA recognition and translational regulation by a homeodomain protein. Nature 379, 694–699.

Fowlkes, C. C., Eckenrode, K. B., Bragdon, M. D., Meyer, M., Wunderlich, Z., Simirenko, L., Luengo Hendriks, C. L., Keranen, S. V., Henriquez, C., Knowles, D. W. et al. (2011). A conserved developmental patterning network produces quantitatively different output in multiple species of Drosophila. PLoS Genet 7, e1002346.

Fowlkes, C. C., Hendriks, C. L., Keranen, S. V., Weber, G. H., Rubel, O., Huang, M. Y., Chatoor, S., DePace, A. H., Simirenko, L., Henriquez, C. et al. (2008). A quantitative spatiotemporal atlas of gene expression in the Drosophila blastoderm. Cell 133, 364–374.

Frasch, M. and Levine, M. (1987). Complementary patterns of even-skipped and fushi tarazu expression involve their differential regulation by a common set of segmentation genes in Drosophila. Genes & development 1, 981–995.

Frohnhöfer, H. G. and Nüsslein-Volhard, C. (1986). Organization of anterior pattern in the Drosophila embryo by the maternal gene bicoid. Nature 324, 120–125.

Fujioka, M., Emi-Sarker, Y., Yusibova, G. L., Goto, T. and Jaynes, J. B. (1999). Analysis of an even-skipped rescue transgene reveals both composite and discrete neuronal and early blastoderm enhancers, and multi-stripe positioning by gap gene repressor gradients. Development (Cambridge, England) 126, 2527–2538.

Goto, T., Macdonald, P. and Maniatis, T. (1989). Early and late periodic patterns of *even skipped* expression are controlled by distinct regulatory elements that respond to different spatial cues. Cell 57, 413–422.

Gursky, V. V., Panok, L., Myasnikova, E. M., Manu, Samsonova, M. G., Reinitz, J. and Samsonov, A. M. (2011). Mechanisms of gap gene expression canalization in the Drosophila blastoderm. BMC Systems Biology 5, 118.

Hengenius, J. B., Gribskov, M., Rundell, A. E., Fowlkes, C. C. and Umulis, D. M. (2011). Analysis of Gap Gene Regulation in a 3D Organism-Scale Model of the Drosophila melanogaster Embryo. PLoS ONE 6, e26797.

Hughes, S. C. and Krause, H. M. (2001). Establishment and maintenance of parasegmental compartments. Development 128, 1109–1118.

Hulskamp, M., Pfeifle, C. and Tautz, D. (1990). A morphogenetic gradient of hunchback protein organizes the expression of the gap genes Kruppel and knirps in the early Drosophila embryo. Nature 346, 577–580.

Jäckle, H., Tautz, D., Schuh, R., Seifert, E. and Lehmann, R. (1986). Cross-regulatory interactions among the gap genes of Drosophila.

Jacob, Y., Sather, S., Martin, J. R. and Ollo, R. (1991). Analysis of Kruppel control elements reveals that localized expression results from the interaction of multiple subelements. Proc Natl Acad Sci U S A 88, 5912–5916.

Jaeger, J., Blagov, M., Kosman, D., Kozlov, K. N., Manu, Myasnikova, E., Surkova, S., Vanario-Alonso, C. E., Samsonova, M., Sharp, D. H., et al. (2004a). Dynamical analysis of regulatory interactions in the gap gene system of Drosophila melanogaster. Genetics 167, 1721–1737.

Jaeger, J. (2011). The gap gene network. Cellular and Molecular Life Sciences 68, 243–274.

Jaeger, J., Manu and Reinitz, J. (2013). Drosophila blastoderm patterning. Current opinion in genetics & development

Jaeger, J., Surkova, S., Blagov, M., Janssens, H., Kosman, D., Kozlov, K. N., Manu, Myasnikova, E., Vanario-Alonso, C. E., Samsonova, M., et al. (2004b). Dynamic control of positional information in the early Drosophila embryo. Nature 430, 368–371.

Janssens, H., Crombach, A., Richard Wotton, K., Cicin-Sain, D., Surkova, S., Lu Lim, C., Samsonova, M., Akam, M. and Jaeger, J. (2013). Lack of tailless leads to an increase in expression variability in Drosophila embryos. Developmental biology

Keränen, S. V. E., Fowlkes, C. C., Luengo Hendriks, C. L., Sudar, D., Knowles, D. W., Malik, J. and Biggin, M. D. (2006). Three-dimensional morphology and gene expression in the Drosophila blastoderm at cellular resolution II: dynamics. Genome Biology 7, R124.

Kozlov, K., Surkova, S., Myasnikova, E., Reinitz, J. and Samsonova, M. (2012). Modeling of gap gene expression in Drosophila Kruppel mutants. PLoS Computational Biology 8, e1002635.

Kraut, R. and Levine, M. (1991a). Mutually repressive interactions between the gap genes giant and Kruppel define middle body regions of the Drosophila embryo. Development 111, 611–621.

Kraut, R. and Levine, M. (1991b). Spatial regulation of the gap gene giant during Drosophila development. Development 111, 601–609.

Lawrence, P. A. (1992). The making of a fly: the genetics of animal design.

Liu, F., Morrison, A. H. and Gregor, T. (2013). Dynamic interpretation of maternal inputs by the Drosophila segmentation gene network. Proceedings of the National Academy of Sciences 110, 6724–6729.

Luengo Hendriks, C. L., Keränen, S. V., Biggin, M. D. and Knowles, D. W. (2007). Automatic channel unmixing for high-throughput quantitative analysis of fluorescence images. Optics express 15, 12306–12317.

Luengo Hendriks, C. L., Keränen, S. V. E., Fowlkes, C. C., Simirenko, L., Weber, G. H., DePace, A. H., Henriquez, C., Kaszuba, D. W., Hamann, B., Eisen, M. B., et al. (2006). Three-dimensional morphology and gene expression in the Drosophila blastoderm at cellular resolution I: data acquisition pipeline. Genome Biology 7, R123.

Manoukian, A. S. and Krause, H. M. (1992). Concentration-dependent activities of the even-skipped protein in Drosophila embryos. Genes Dev 6, 1740–1751.

Manu, Surkova S., Spirov, A. V., Gursky, V. V., Janssens, H., Kim, A.-R., Radulescu, O., Vanario-Alonso, C. E., Sharp, D. H., Samsonova, M., et al. (2009a). Canalization of gene expression and domain shifts in the Drosophila blastoderm by dynamical attractors. PLoS Computational Biology 5, e1000303.

Manu, Surkova S., Spirov, A. V., Gursky, V. V., Janssens, H., Kim, A.-R., Radulescu, O., Vanario-Alonso, C. E., Sharp, D. H., Samsonova, M., et al. (2009b). Canalization of gene expression in the Drosophila blastoderm by gap gene cross regulation. PLoS biology 7, e1000049.

Martinez-Arias, A. and Lawrence, P. A. (1985). Parasegments and compartments in the Drosophila embryo. Nature 313, 639–642.

Mohr, S. E. and Perrimon, N. (2012). RNAi screening: new approaches, understandings, and organisms. Wiley Interdiscip Rev RNA 3, 145–158.

Namba, R., Pazdera, T. M., Cerrone, R. L. and Minden, J. S. (1997). Drosophila embryonic pattern repair: how embryos respond to bicoid dosage alteration. Development 124, 1393–1403.

Nasiadka, A. and Krause, H. M. (1999). Kinetic analysis of segmentation gene interactions in Drosophila embryos. Development 126, 1515–1526.

Ni, J. Q., Zhou, R., Czech, B., Liu, L. P., Holderbaum, L., Yang-Zhou, D., Shim, H. S., Tao, R., Handler, D., Karpowicz, P. et al. (2011). A genome-scale shRNA resource for transgenic RNAi in Drosophila. Nat Methods 8, 405–407.

Nüsslein-Volhard, C., Frohnhöfer, H. G. and Lehmann, R. (1987). Determination of anteroposterior polarity in Drosophila. Science 238, 1675–1681.

Ochoa-Espinosa, A., Yucel, G., Kaplan, L., Pare, A., Pura, N., Oberstein, A. L., Papatsenko, D. and Small, S. (2005). The role of binding site cluster strength in Bicoid-dependent patterning in Drosophila. Proceedings of the National Academy of Sciences of the United States of America 102, 4960–4965.

Papatsenko, D. and Levine, M. S. (2011). The Drosophila gap gene network is composed of two parallel toggle switches. PLoS ONE 6, e21145.

Pignoni, F., Steingrímsson, E. and Lengyel, J. A. (1992). bicoid and the terminal system activate tailless expression in the early Drosophila embryo. Development (Cambridge, England) 115, 239–251.

Pisarev, A., Poustelnikova, E., Samsonova, M. and Reinitz, J. (2009). FlyEx, the quantitative atlas on segmentation gene expression at cellular resolution. Nucleic Acids Res 37, D560–D566.

Rivera-Pomar, R., Lu, X., Perrimon, N., Taubert, H. and Jackle, H. (1995). Activation of posterior gap gene expression in the Drosophila blastoderm. Nature 376, 253–256.

Saulier-Le Drean, B., Nasiadka, A., Dong, J. and Krause, H. M. (1998). Dynamic changes in the functions of Odd-skipped during early Drosophila embryogenesis. Development 125, 4851–4861.

Schroeder, M. D., Greer, C. and Gaul, U. (2011). How to make stripes: deciphering the transition from non-periodic to periodic patterns in Drosophila segmentation. Development (Cambridge, England) 138, 3067–3078.

Small, S., Blair, A. and Levine, M. S. (1996). Regulation of two pair-rule stripes by a single enhancer in the Drosophila embryo. Developmental biology 175, 314–324.

Small, S., Kraut, R., Hoey, T., Warrior, R. and Levine, M. S. (1991). Transcriptional regulation of a pair-rule stripe in Drosophila. Genes & Development 5, 827–839.

St Johnston, D. and Nüsslein-Volhard, C. (1992). The origin of pattern and polarity in the Drosophila embryo. Cell 68, 201–219.

Staller, M. V., Yan, D., Randklev, S., Bragdon, M. D., Wunderlich, Z. B., Tao, R., Perkins, L. A., Depace, A. H. and Perrimon, N. (2013). Depleting gene activities in early Drosophila embryos with the “maternal-Gal4-shRNA” system. Genetics 193, 51–61.

Stern, D. L., and E. Sucena. (2000). Preparation of larval and adult cuticles for light microscopy. In Drosophila protocols pp. 601–615.

Struhl, G., Struhl, K. and Macdonald, P. M. (1989). The gradient morphogen bicoid is a concentration-dependent transcriptional activator. Cell 57, 1259–1273.

Surkova, S., Golubkova, E., Manu, Panok, L., Mamon, L., Reinitz, J. and Samsonova, M. (2013). Quantitative dynamics and increased variability of segmentation gene expression in the Drosophila Krüppel and knirps mutants. Developmental biology 376, 99–112.

Tamari, Z. and Barkai, N. (2012). Improved readout precision of the Bicoid morphogen gradient by early decoding. J Biol Phys 38, 317–329.

Tautz, D. (1988). Regulation of the Drosophila segmentation gene hunchback by two maternal morphogenetic centres. Nature 332, 281–284.

Waddington, C. H. (1942). Canalization of Development and the inheritance of acquired characters. Nature 563–565.

Werz, C., Lee, T. V., Lee, P. L., Lackey, M., Bolduc, C., Stein, D. S. and Bergmann, A. (2005). Mis-specified cells die by an active gene-directed process, and inhibition of this death results in cell fate transformation in Drosophila. Development 132, 5343–5352.

Wieschaus, E., Nusslein-Volhard, C. and Kluding, H. (1984). Kruppel, a gene whose activity is required early in the zygotic genome for normal embryonic segmentation. Dev Biol 104, 172–186.

Wunderlich, Z., Bragdon, M. D., Eckenrode, K. B., Lydiard-Martin, T., Pearl-Waserman, S. and DePace, A. H. (2012). Dissecting sources of quantitative gene expression pattern divergence between Drosophila species. Mol Syst Biol 8, 604.

